# Theory of local k-mer selection with applications to long-read alignment

**DOI:** 10.1101/2021.05.22.445262

**Authors:** Jim Shaw, Yun William Yu

## Abstract

**Motivation:** Selecting a subset of k-mers in a string in a local manner is a common task in bioinformatics tools for speeding up computation. Arguably the most well-known and common method is the minimizer technique, which selects the ‘lowest-ordered’ k-mer in a sliding window. Recently, it has been shown that minimizers are a sub-optimal method for selecting subsets of k-mers when mutations are present. There is however a lack of understanding behind the theory of why certain methods perform well.

**Results:** We first theoretically investigate the conservation metric for k-mer selection methods. We derive an exact expression for calculating the conservation of a k-mer selection method. This turns out to be tractable enough for us to prove closed-form expressions for a variety of methods, including (open and closed) syncmers, (*α, b, n*)-words, and an upper bound for minimizers. As a demonstration of our results, we modified the minimap2 read aligner to use a more optimal k-mer selection method and demonstrate that there is up to an 8.2% relative increase in number of mapped reads.

**Availability and supplementary information:** Simulations and supplementary methods available at https://github.com/bluenote-1577/local-kmer-selection-results. os-minimap2 is a modified version of minimap2 and available at https://github.com/bluenote-1577/os-minimap2.

## 1 Introduction

In recent decades, there has been an exponential increase in the amount and throughput of available sequencing data (Goodwin *et al.*, 2016; Schmidt and Hildebrandt, 2017), necessitating more efficient modern methods for processing sequencing data (Berger *et al.*, 2016). Many methods employ *k-mer* (length-k substrings of a sequence) based analysis, because k-mer methods tend to be fast and memory efficient. k-mer methods appear in metagenomics (Wood and Salzberg, 2014), genome assembly (Nagarajan and Pop, 2013; Rautiainen and Marschall, 2020), read alignment (Li, 2018), variant detection (Peterlongo *et al.*, 2017; Shajii *et al.*, 2016), transcriptomics (Bray *et al.*, 2016), and many more. Because k-mers overlap, selecting a subset of k-mers in a sequence can for many applications lead to a dramatic increase in efficiency while only losing a small amount of information. In this paper, we will focus on *local* k-mer selection methods, which means that the criteria for selecting a specific k-mer should depend on the local information near the k-mer.

A popular class of local selection methods revolve around *minimizers* (Schleimer, 2003; Roberts *et al.*, 2004), and there is a lot of recent literature on both practically optimizing minimizer efficiency and theoretical intrinsic properties of minimizers (Marçais *et al.*, 2018; Zheng *et al.*, 2021, 2020a,b). Recently, new local selection methods for k-mer selection have been proposed which are distinct from minimizers, including *syncmers* (Edgar, 2021) and *minimally-overlapping words* (Frith *et al.*, 2020).

Improvements for minimizer techniques have historically focused on optimizing for *density*, the fraction of selected k-mers. However, it often makes more sense for density to be an application dependent tunable parameter. Thus, Edgar (2021) instead propose the new metric of *conservation,* which measures the fraction of bases in a sequence which can be “recovered” by k-mer matching after the sequence undergoes a random mutation process; a similar metric is also used in Frith *et al.* (2020) and Sahlin (2021). While newer techniques have demonstrated effectiveness through empirical studies, it is not clear *why* certain methods perform well beyond heuristic notions.

### 1.1 Contributions

We make both theoretical and practical contributions in this manuscript. The first part of our paper is theoretical. In Sections 2,3 and 4 we develop a novel, more general, mathematical framework for analyzing local k-mer selection methods. We show how our framework rigorously discerns the relationship between the notion of “clumping” or “overlapping” of k-mers alluded to in Edgar (2021); Frith *et al.* (2020) and conservation (Theorem 2). We then mathematically analyze existing local k-mer selection methods, resulting in new closed-form expressions of conservation for various k-mer selection methods and a novel result on optimal parameter choice for open syncmers (Theorem 6).

The second part of our contributions is practical. In Section 5, we empirically calculate conservation for a wide range of methods for which we can not derive a closed-form expression. We then modified the existing software minimap2 (Li, 2018) to use open syncmers, which we found to be a well-conserved k-mer selection method, instead of minimizers. Our results show that 1) conservation and alignment sensitivity are correlated and that 2) alignment sensitivity is increased after modifying minimap2. This is, to our knowledge, the first experiment showing that practical software for long-read alignment can be improved by considering better k-mer selection methods.

## 2 Preliminaries

We formally define what a local k-mer selection method is in this section. We give a general formalism that is original and extends the existing formalisms in Marçais *et al.* (2018). We also review existing local k-mer selection methods.

### 2.1 k-mer selection methods

Let Σ be our alphabet. We will be implicitly dealing with nucleotides (Σ = {*A, C, G, T*}) for the rest of the paper, although our results generalize without issues. For a string *S* ∈ Σ*, we use the notation *S*[*i, k*] to mean the substring of length k starting at index *i*. We will assume our strings are 1-indexed.

#### Definition 1.

*A k-mer selection method is a function f from the set of finite strings* Σ* *such that for S ∈* Σ*, *f*(*S*) *contains (possibly multiple) tuples* (*x, i*) *where x* ∈ Σ^*k*^ *is a k-mer in S, and i is the starting position where x occurs*.

We will sometimes refer to a k-mer selection method as a selection method or just a method when the value of *k* is implied. We now define a *local* k-mer selection method.

#### Definition 2.

*A method f is a q-local method if*

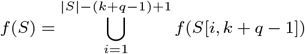

*for every S* ∈ Σ * *of length* ≥ *k* + *q* – 1 *after an appropriate shift in the position of the k-mers*.

In other words, a q-local method is just defined on (*k* + *q* – 1)-mers and then extended to arbitrary strings. The special case of *q* = 1 implies that the method can be defined by examining all k-mers and deciding if each k-mer is selected or not. We will always assume that |*S*| ≥ *q* + *k* – 1, and will focus only on local methods in this paper. The main reason for doing so is that q-local methods have the following desirable property that is an easy result of the definition.

#### Theorem 1.

*Let f be a q-local method. If two strings S, S′ share a region of length k* + *q* – 1, *i.e. S*[*i, k* + *q* – 1] = *S′* [*j, k* + *q* – 1], *then every k-mer in f* (*S*[*i, k* + *q* – 1]) = *f* (*S′* [*j, k* + *q* – 1]) *is also in f* (*S′*) *and f* (*S*) (*if ignoring the index of the starting position*).

Proof. Follows easily from the definition of q-locality.

To see not all k-mer selection methods are local, it is not hard to see that the MinHash (Ondov *et al.*, 2016; Broder, 1998) sketch, which is computed by selecting a fixed number of k-mers hashing to the smallest values over the *entire* genome, is not local. Another method that is not local is selecting every *n*th k-mer occurring in a string because it depends on the global property of starting position. In the next section, we will give examples of local methods.

Our notion of a local k-mer selection method is more general than the notion of local schemes defined in Marçais *et al.* (2018). Local schemes are defined to be functions of the form

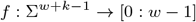

where [0: *w* – 1] = {0,…, *w* – 1}. Local schemes essentially select exactly one k-mer from a *w* + *k* – 1-mer by specifying the starting location for a specific k-mer. While local schemes give rise to *w*-local k-mer selection methods, not all local k-mer selection methods are local schemes because local selection methods may select 0 or more than 1 k-mer in a window. Local schemes are defined in such a way to satisfy a property called the *window guarantee*.

#### Definition 3.

*A local k-mer selection method has the r-window guarantee property for r if* 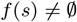 *for all s* ∈ Σ^*k*+*r*−1^.

The window guarantee says that for every *r* consecutive k-mers, the local method will select at least one k-mer, guaranteeing that there will be no large gaps on the string for which no k-mer is selected. While the window guarantee is useful for many applications (Marçais *et al.*, 2019), in some applications such as alignment it is not necessary. A closely related notion to the window guarantee is universal hitting sets (Ekim *et al.*, 2020; Orenstein *et al.*, 2017), which give rise to local methods with a window guarantee. However, we will not discuss such methods because they are not computationally efficient; e.g. PASHA (Ekim *et al.*, 2020) takes > 95 hours to compute a UHS for *k* =16.

Another important property of a selection method we will discuss is the *density*.

#### Definition 4.

*The density of a k-mer selection method f is the expected number of selected k-mers divided by the number of total k-mers in a uniformly random string with i.i.d characters S as* |*S*| → ∞.

Given a long uniform random string *S*, by taking indicator random variables *Y_i_* for the event that the k-mer starting at *i* is in *f*(*S*), one can easily see density is equal to Pr(*Y_i_* = 1) as long as *i* is not near the edges (the start/end) of *S*. The issue near the edges is that in Definition 2, a k-mer near the start of *S* appears in less of the *S*[*i, k* + *q* – 1] “windows” than a k-mer in the middle of the string. This is known as the edge bias problem in minimizers (Edgar, 2021).

### 2.2 Overview of specific selection methods

There are three overarching classes of local selection methods we will define in this section. These methods have been discussed previously in Zheng *et al.* (2020a); Frith *et al.* (2020); Edgar (2021); Orenstein *et al.* (2017) but we will give a brief review and outline the specific properties associated to each method.

#### 2.2.1 Minimizers

The most well-known and frequently used class of local k-mer selection methods are minimizer methods, originally appearing in Roberts *et al.* (2004); Schleimer (2003).

##### Definition 5.

*Given a triple* 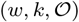 *where w, k are integers and* 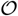 *is an ordering on the set of all k-mers, a minimizer outputs the smallest k-mer appearing in a* (*w* + *k* – 1)-*mer, or equivalently a window of w consecutive k-mers*.

From now on, when we specify the smallest value in a window, ties are broken by letting the leftmost k-mer be the smallest. A minimizer gives rise to a *w*-local method with a *w*-window guarantee by examining all windows of *w* k-mers and selecting k-mers inside this window. *k* and *w* are application dependent parameters that control for density and k-mer size, so there is one free parameter which is the ordering 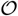.

Somewhat surprisingly, the ordering, which can be string-dependent, plays a very important part of minimizer performance. We define a *random minimizer* to be a minimizer with the ordering defined by a random permutation. One can show that the density for a random minimizer is 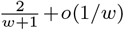 (Marçais *et al.*, 2018) for “reasonable” *w* and *k.* However, with a specific ordering, one can achieve provably better densities for the same parameters. For example, the miniception (Zheng *et al.*, 2020a) algorithm gives a density upper bounded by 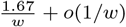.

##### Definition 6

(Charged contexts). *Given parameters* 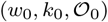, *a window of consecutive w*_0_ + 1 *k*_0_-*mers (equivalently a* (*w*_0_ + *k*_0_)-*mer) is a charged context if the smallest k*_0_-*mer is the first or last k*_0_-*mer in the window*.

##### Definition 7

(Miniception).

- *Let w, k, k*_0_ *be integers, and define w*_0_ = *k* – *k*_0_. *Define a random minimizer* 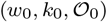.
- *Let C*_0_ *be the set of charged contexts for* 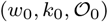. *Note that each element of C*_0_ *is a k-mer*.
- *Define a new minimizer* 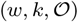 *where the k-mers in C*_0_ *are less than all other k-mers not in C*_0_ *and with random orderings in C*_0_ *and* Σ^*k*^ \ *C*_0_.

There is a subtlety in that *k*_0_ gives a free parameter. It is not obvious how to set this parameter; a heuristic is given in Zheng *et al.* (2020a). In this paper, we will only discuss random minimizers and the miniception.

#### 2.2.2 Syncmers

Syncmers were first defined in Edgar (2021) to be a class of 1-local k-mer selection methods. These methods break k-mers into a window of *k* – *s* + 1 consecutive s-mers, for some parameter *s* < *k*, and select the k-mer based on some criteria involving the smallest s-mers in the window. From now on, we will assume the ordering on the s-mers is random.

We have already seen such a construction; it was pointed out in Edgar (2021) that the charged contexts described in the definition of the miniception (Definition 6) are also called *closed syncmers* in the terminology of Edgar (2021), where *k*_0_ = *s* and *w*_0_ = *k* – *s*. If one simply takes the set of closed syncmers to be the selected k-mers instead of using them as “seeds” for the miniception, that would be a valid 1-local k-mer selection method. Closed syncmers can be shown to have a *k* – *s* window guarantee, so this is our first (and only) example of a method that is 1-local and has a window guarantee.

While there are variations on syncmer methods, we will primarily analyze the *open syncmer* defined in Edgar (2021) due to it being shown empirically to be the best performing with respect to the conservation metric, which we will discuss in Section 3.

##### Definition 8

(Open syncmer). *Let* (*k, s, t*) *be parameters with s < k and* 1 ≤ *t* ≤ *k* – *s* + 1. *Considering a k-mer as a window of k – s* + 1 *consecutive s-mers, a k-mer is selected if the smallest s-mer appears at position t in the window*.

Importantly, open syncmers *do not* have a window guarantee window guarantee. Consider the string AAAAA… with *t* = 2 for any *k, s*. Remembering that the leftmost s-mer is the smallest by convention, the smallest s-mer always occurs at the first position so no k-mers are selected by this method.

The density of open syncmers and closed syncmers are respectively 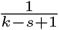 and 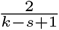 up to a small 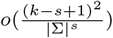 error term, following the exact same argument as Edgar (2021) and Section 2.3.1 of Zheng *et al.* (2020a) for the error term.

#### 2.2.3 Words based method

Another class of 1-local selection methods is to consider a set of words *W* ⊂ Σ* and select a k-mer if a prefix of the k-mer lies in *W*. The approach taken in Frith *et al.* (2020) is to consider possible *W*s and find “good” possible choices for *W*. The intuition is that they want the words in *W* to not overlap with each other; this way selected k-mers overlap less. One metric they consider for goodness of a candidate *W* is the variance-to-mean ratio of the number of k-mers appearing in a random string for the method associated to *W*.

They offer a simple form for *W* by letting *W* = {*x* ∈ Σ^*n* + 1^: *x* = *abbb*…} where *a* = *A* and *b* can be any of{*C, T, G*} for the nucleotide alphabet. We will refer to this as the (*a, b, n*)-words method. The density of this method is 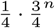.

More sophisticated choices for *W* can be constructed. We will not analyze such methods theoretically since these *W* are found by an optimization algorithm and thus hard to analyze. We will instead test against more sophisticated *W* empirically.

**Table 1.**
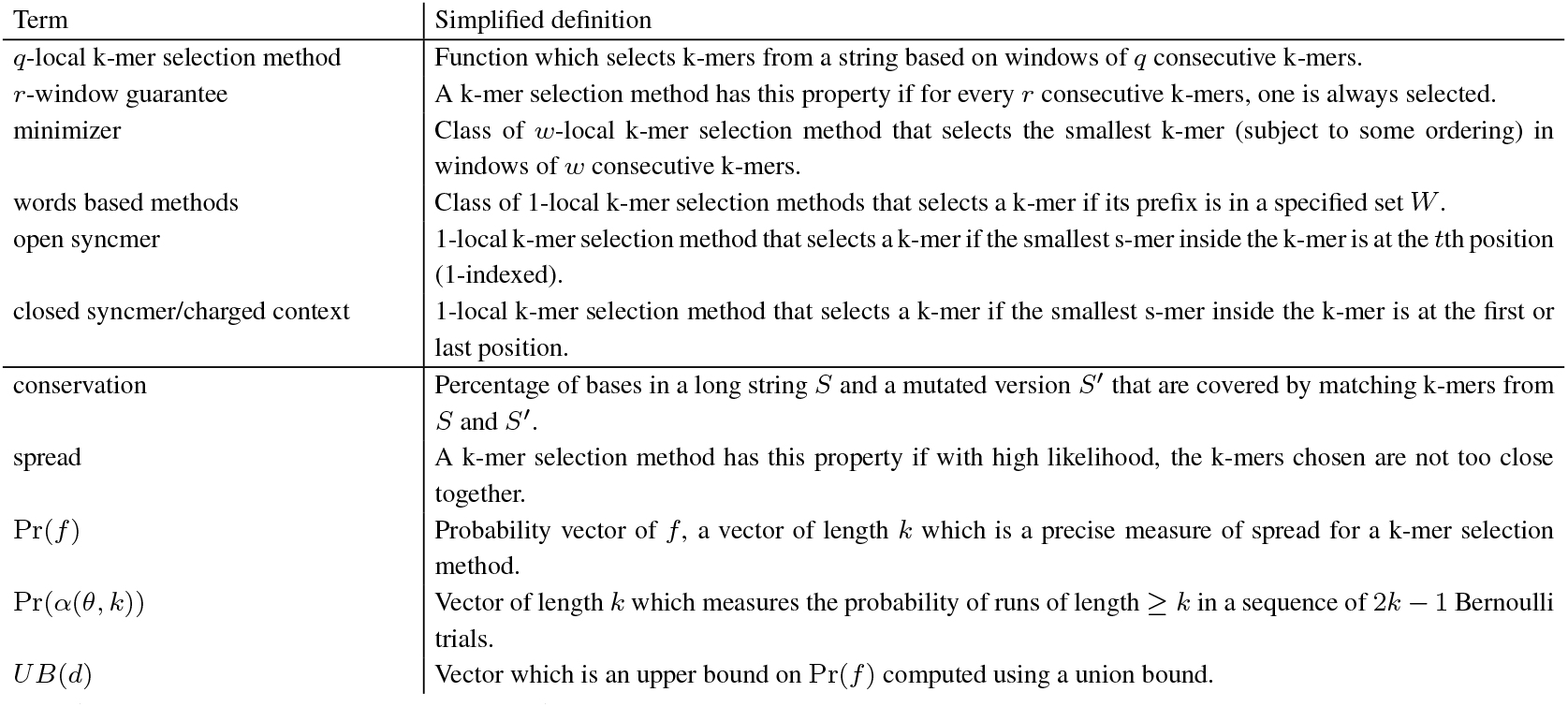
Simplified definitions of concepts discussed in Sections 2 and 3.

**Table 2.**
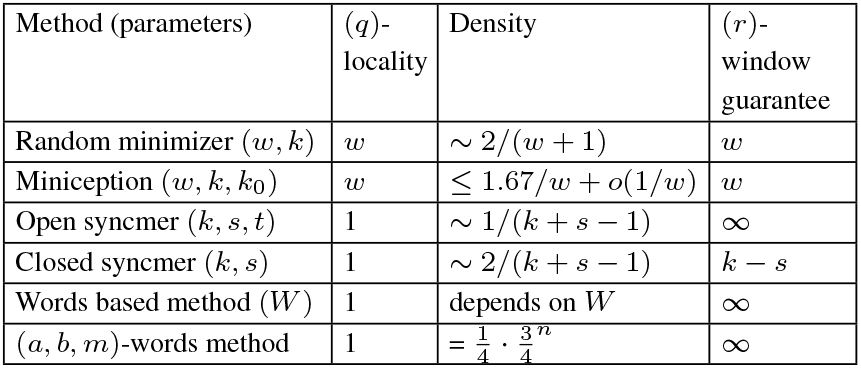
Properties of discussed methods. The ~ sign denotes up to a small error term. ∞ means that there is no window guarantee.

## 3 Analytical framework

While we have discussed intrinsic features of selection methods such as locality, density, and window guarantee, we have not yet discussed the “performance” of selection methods. It turns out that evaluating the performance of a method is subtle and depends on the task at hand. In certain contexts, methods can not be compared because some tasks such as constructing de Bruijn graphs from minimizers (Rautiainen and Marschall, 2020) or counting and binning k-mers (Marçais *et al.*, 2019) require a window guarantee while other tasks such as alignment do not.

To compare methods, we will fix the density across methods and evaluate a precise notion of performance which we will define below. In previous studies on minimizers (Marçais *et al.*, 2018; Zheng *et al.*, 2020a; Marçais *et al.*, 2017) the focus was only on optimizing the density for a fixed window size w, the assumption being that a method’s performance is *only reliant on the size of the window guarantee*. This is not an unreasonable assumption for certain tasks, but it is not applicable to methods without a window guarantee.

When selecting a subset of k-mers, a good method should select k-mers that are spread apart on *S*, i.e. k-mers should not overlap or clump together. This is noted in Frith *et al.* (2020). Intuitively, this is because close together k-mers give similar information due to more base overlaps. To formalize this, in Section 3.1 we give a new, precise notion of spread. In Section 3.3, we prove an original result detailing how this formalism relates to conservation.

### 3.1 Formalizing k-mer spread

Let *S* = *x*_1_*x*_2_… be a long random string of fixed length with independent and uniformly random characters over Σ. We now define a key quantity associated to *f* which we will call the probability vector of *f*.

#### Definition 9

(Probability vector of *f*). *Let i be a position in S which is away from the edges. Define the event E_j_* = {(*S*[*j, k*],*j*) ∈ *f*(*S*)} *representing whether or not a k-mer at position j is selected by f, and*

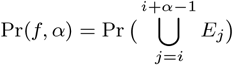

*the probability that some k-mer is selected from S*[*i,k* + *α* – 1]. *We call*

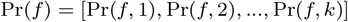

*the probability vector of f*.

This notion is well-defined because our string consists of i.i.d letters, and our method *f* is translation invariant along the string due to locality. As long as *i* is not near the end or beginning of *S*, the choice of *i* does not matter. Notice that Pr(*f*, 1) = Pr(*E_i_*) = Pr(*E_j_*) for any *j* is the probability that a random k-mer is selected by *f*, which is the density of *f*. Therefore Pr(*f, α*) measures how disjoint the sets *E_i_*,…, *E*_*i*+*α*−1_ are. The more disjoint the events *E_i_*,…, *E*_*i*+*α*−1_ are the more spread out the selected k-mers are.

We can get a natural upper bound for Pr(*f*) because 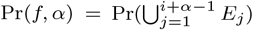 so the union bound gives

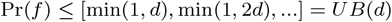

where ≤ means over all components, remembering that Pr(*E_j_*) = *d* is the density. Asymptotically, the upper bound can be reached. In Marçais *et al.* (2018), it was shown that asymptotically in *k*, one can construct a minimizer with density 1/*w* and window guarantee *w*. This implies that Pr(*f*, 1) = 1/*w* and Pr(*f, w*) = 1, showing that Pr(*f, n*) = *n/w* for all *n* ≤ *w*.

On a technical note, one can actually see that for 1-local methods, Pr(*f, α*) is equivalent to Pr(*f* 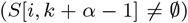. The issue is that for *w*-local methods, *f*(*S*[*i, k* + *α* – 1]) is not defined if *α* < *w*. For example, one can not deduce if a k-mer is selected by a minimizer method just based on the k-mer itself; a window of k-mers is needed.

### 3.2 Mutated k-mer model

Let 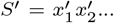 be a mutated version of *S* such that 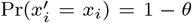 and 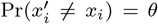, where the mutated character is uniform over the rest of the alphabet and *θ* ∈ [0, 1] is some mutation parameter. A similar model for k-mer mutations is used in Blanca *et al.* (2021). We now give a mathematical definition of the conservation metric which is used in Edgar (2021).

#### Definition 10

(Conservation). *Given a k-mer selection method f and parameter θ, let the set of conserved bases be*

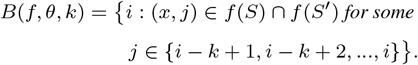

*Define the conservation to be* 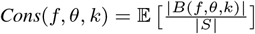.

The set *B*(*f, θ, k*) is the set of bases for which 1) a k-mer is selected by *f* overlapping the base and 2) this k-mer is *unmutated* from *S* to *S*′. Notice in our definition that for a position to be covered, we require the position of the k-mers to be the same in *S* and *S*′ so we disregard spurious matches across the genome.

### 3.3 Relating conservation and spread

We now show that Cons(*f, θ, k*) and Pr(*f*), which captures k-mer spread, are related. To calculate Cons(*f, θ, k*), we let *X_i_* be indicator random variables where *X_i_* = 1 if *i* ∈ *B*(*f, θ, k*). By linearity of expectation

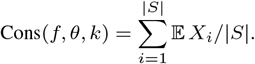

The *X_i_* are not independent; if *i* ∈ *B*(*f, θ,k*), then it is likely that *i* + 1 ∈ *B*(*f, θ, k*) as well. If *i* ≠ *j* and both lie away from the ends of *S*, we get that Pr(*X_i_* = 1) = Pr(*X_j_* = 1). If *k* ≪ |*S*| the contribution from positions near at the edges of the string is small. Therefore, we will make the assumption 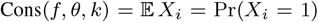 for some *i* in *S* away from the edges.

#### 3.3.1 Understanding mutation configurations

Given the base *i,* the k-mers covering *i* on *S*′ are *S*′ [*i* – *k* + 1, *k*],…, *S*′ [*i, k*] so the substring of all covered bases is *S*′ [*i* – *k* +1, 2*k* – 1]. We examine how mutations change the k-mers for these bases. We can consider these 2*k* – 1 bases on *S*′ as 2*k* – 1 Bernoulli trials with success probability 1 – *θ* corresponding to an unchanged base. We call possible sequences of Bernoulli trials configurations of the mutations. A similar notion of configurations is used in Chaisson and Tesler (2012).

Given some configuration, we will get *α* unmutated k-mers overlapping *i*, where *α* ∈ {0,…, *k*}, an unmutated k-mer being one for which all bases are unmutated. These unmutated k-mers are candidates for being in *f*(*S*) ⋂ *f*(*S*′). In Figure 1 we show graphically how different configurations lead to a different value of *α*.

**Fig. 1:**
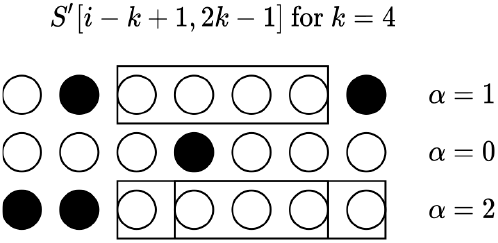
Three examples of mutation configurations and corresponding *α* values for *k* = 4. Circles represent bases of *S*′ around *i*, while black and white indicate mutated or unmutated bases respectively. Boxes indicate an unmutated k-mer overlapping position *i*.

From Figure 1, it is clear that *α* = *α*(*θ, k*) is a random variable corresponding to the number of successful runs of length exactly *k* in 2*k* – 1 trials, or equivalently *α*(*θ, k*) = max{0, *L* – *k* +1} where *L* is the longest run. This is a well-studied problem; see Chapter 5.3 in Uspensky (1965) for a closed-form solution for the probability of a run of length at least *m* in *n* trials.

#### 3.3.2 Calculating 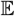 *X_i_* by conditioning

Conditioning on *α*, we get

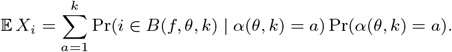

Notice that Pr(*i* ∈ *B*(*f, θ, k*) |*α*(*θ, k*) = *a*) is the probability that some k-mer is selected from the *a* consecutive unmutated k-mers in *S* and *S*′. In the language of Definition 9, letting 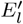 be the same event as *E_l_* but over *S*′, we get

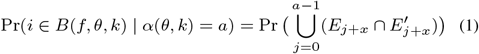

where *x* is arbitrary and can be thought of as the starting position of the first unmutated k-mer (the probability does not depend on *x*). Locality comes into play now; if *f* is a 1-local method, then the event *E*_*j*+*x*_ is true if and only if 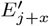 is true. This follows because *S*′ [*j* + *x, k*] = *S*[*j* + *x, k*] by the assumption, and by 1-locality, this k-mer is in *f*(*S*) ⋂ *f*(*S*′) after reindexing if and only if this k-mer is selected by *f*. Hence 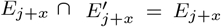 for 1-local methods. Now we see that the right-hand side of Equation 1 is exactly Pr(*f, α*) from Definition 9, resulting in the following theorem.

##### Theorem 2.

*Let* Pr(*α*(*θ,k*)) = [Pr(*α*(*θ, k*) = 1), Pr(*α*(*θ, k*) = 2),… Pr(*α*(*θ, k*) = *k*)] *and f be a 1-local method. Then*

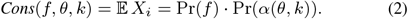

*If f is not 1-local, then*

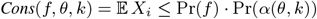

*follows from the trivial upper bound* 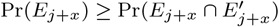.

If *f* is not 1-local, then 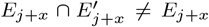. For minimizers, this is the *context dependency problem*; if a k-mer is selected in *S* and the same k-mer is also in *S*′, it may not be selected due to mutations in the window. This shows why 1-local methods are inherently superior for tasks that require k-mer matching for mutated strings such as alignment. We will discuss this discrepancy for random minimizers in more detail in Section 4.2 and show in simulations the extent of this discrepancy.

#### 3.3.3 Calculating Pr(*α*(*θ, k*))

Even though the successful runs problem is solved for general parameters, the formula for the probability is derived by manipulating generating functions and is a bit unruly. For the case of 2*k* – 1 trials and ≥ *k* runs, the derivation can actually be done in a much more straightforward way. The following theorem is proved in Suppelementary Section S1, where we also show plots for Pr(*α*(*θ, k*)) over varying parameters.

##### Theorem 3.

*For* 2*k* – 1 *i.i.d Bernoulli trials with success probability* 1 – *θ and* 0 ≤ *β* ≤ *k* – 1,

Pr(*α*(*θ; k*) = *β* + 1) = Pr(*Longest run of successes is k* + *β*)

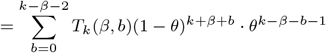

*where*

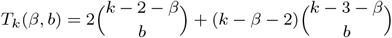

*and binomial coefficients with negative parameters are* 0. *For β* = *k* – 1, *the probability of* 2*k* – 1 *successes is just* (1 – *θ*)^2*k*–1^.

## 4. Mathematical analysis of specific methods

### 4.1 Syncmers

Let *k, s, t* be given and *f* be either a closed or open syncmer method. To calculate Pr(*f, α*), we need to analyze the *α* consecutive k-mers. Breaking these k-mers into s-mers, we get a window of *k* – *s* + *α* s-mers. We will now make an assumption that all s-mers in the window are distinct. For uniform random strings, as shown in the proof of Lemma 9 in Zheng *et al.* (2020a), the probability of two identical s-mers appearing in a window is upper bounded by 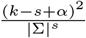. For minimap2 (Li, 2018), the default parameters are *k* = 15 and *w* = 10. To achieve approximately the same density using an open syncmer, we would require *s* = 10. An easy calculation shows that this probability is very small in this regime.

Since all s-mers are distinct and we assume some random ordering on all s-mers, the relative ordering of s-mers in this window is a uniformly random permutation in *σ* ∈ *S*_*k*−*s*+*α*_ where *σ*(*i*) is the relative ordering of position *i* in the window. Determining whether or not a k-mer is selected then amounts to analyzing a random permutation’s smallest elements; see Figure 2.

**Fig. 2:**
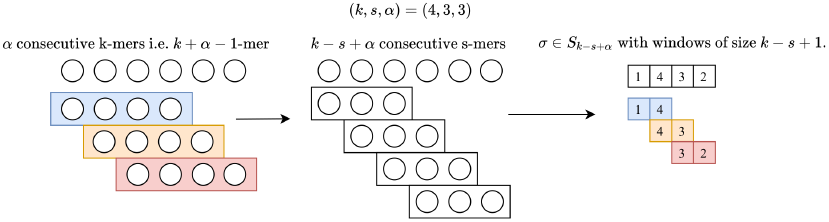
A graphical example of how *α* consecutive k-mers gives rise to a permutation in *S*_*k*–*s*+*α*_ where the first s-mer is the smallest, the second is the largest, and so forth. The colors show how k-mers correspond to windows in the permutation.

Now given a permutation *σ* ∈ *S*_*k*–*s*+*α*_, we consider all windows [*σ*(*i*), *σ*(*i* + 1),…, *σ*(*i* + *k* – *s*)] of size *k* – *s* + 1 corresponding to the s-mers inside a k-mer starting at position *i*. The permutation is “successful”, i.e corresponds to the event that some k-mer is chosen by *f*, if one of these k-mers is chosen by *f*. We now count the number of successful permutation for open syncmers and closed syncmers.

#### Theorem 4

(Successful permutations for closed syncmers). *Let CS*(*α, k, s*) *be the number of permutations in S*_*k*–*s*+*α*_ *such that for some window* [*σ*(*i*),…, *σ*(*i* + *k* – *s*)], *either σ*(*i*) *or σ*(*i* + *k* – *s*) *is the smallest element in the window. If α* ≤ *k* – *s*,

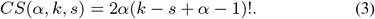

*If α* > *k* – *s, then CS*(*α, k, s*) = (*k* ‒ *s* + *α*)!.

#### Corollary 1.

*If f is a closed syncmer method, then*

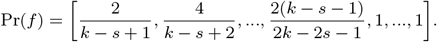

The corollary follows by seeing that Pr(*f, α*) = *CS*(*α,k,s*)/(*k* – *s* + *α*)!, which comes from our discussion about how random consecutive k-mers give rise to uniformly random permutations under our assumptions. Notice that Pr(*f*, 1) = 2/(*k* ‒ *s* +1) and Pr(*f, k* – *s*) = 1 which is in line with the density and window guarantee discussed in Table 2.

#### Proof of Theorem 4.

Let *σ* ∈ *S*_*k*–*s*+*α*_. Note that *α* is the number of windows in *σ* of size *k* – *s* + 1. For *i* ∈ *A_α_* = {1,…, *α*}, if *σ*(*i*) = 1 then the first s-mer in some window starting at *i* is the smallest, so this permutation is successful. Otherwise, for *i* ∈ *B_α_* = {*k* – *s* + 1,…, *k* – *s* + *α*} if *σ*(*i*) = 1 then the last s-mer in some window is the smallest, so it is successful.

Suppose *σ*(i) ≠ 1 for all *i* ∈ *B_α_* ∪ *A_α_*, so *σ*(*i*) = 1 for some *i* ∈ {*α* + 1,…, *k* – *s*}. Notice that every single window contains all positions {*α* + 1,…, *k* – *s*}, but these positions are neither the start nor end of the window. Therefore *σ* is successful if and only if *σ*(*i*) = 1 for some *i* ∈ *B_α_* ∪ *A_α_*. Counting permutations gives 2*α*(*k* – *s* + *α* – 1)! possible permutations.

We can also count open syncmers. Unfortunately, the number of permutations is only determined as a recurrence relation and the formula is not as nice. Theorem 5 is proved in Supplementary Section S2.

#### Theorem 5.

*Successful permutations for open syncmers] Using parameters k, s, t as defined in Definition* 8, *let τ* = *t* – 1 *and OS*(*α, k, s, t*) *be the number of permutations in S*_*k*–*s*+*α*_ *such that for some window* [*σ*(*i*),…, *σ*(*i* + *k* – *s*)] *the smallest element is σ*(*i* + *τ*).

*Define ℓ*_1_ = *τ*, *ℓ*_2_ = *k* – *s* – *τ*. Then

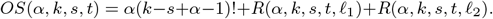

*We define R*(*α, k, s, t, ℓ*) *as*

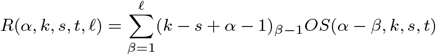

*where the subscript indicates falling factorial, and OS*(*α*–*β, k, s, t*) = 0 *if β* ≥ *α*.

One can divide *OS*(*α, k, s, t*) by (*k* – *s* + *α*)! to get Pr(*f, α*) as previously discussed. This formula does not seem to have a nice closed-form, but it still allows us to answer a question about the optimal choice of parameter *t*. In Edgar (2021), the impact of parameter *t* was investigated on the performance of open syncmers, but no explicit conclusion was made about how to choose *t*. Below we prove a theorem that says that the parameter *t* should be chosen to be the middle position of a window of *k* – *s* + 1 s-mers.

#### Theorem 6.

*Let* 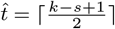. *Then* 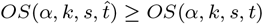 *for any valid choice of t*.

We also prove Theorem 6 in Supplementary Section S2. Note that *t* ↦ *k* – *s* + 2 – *t* gives the same Pr(*f*) by Theorem 5. By Theorems 2 and 6, we now have a rigorous justification of the optimal value for t to maximize conservation. Thus, *t* is not actually a free parameter when optimizing for conservation.

### 4.2 Random minimizer

Let *w, k* be parameters for a random minimizer method *f*. To calculate Pr(*f, α*), as opposed to the analysis in the syncmer section where we look only at *α* consecutive k-mers, we now have to look *all* k-mers in all windows containing any of these *α* consecutive k-mers. As in the syncmer case, we will assume all k-mers are distinct.

Given *α* consecutive k-mers, we need to also know the ordering of the *w* – 1 k-mers to the left and to the right of these *α* k-mers because they are included in some window containing one of these consecutive k-mers. This gives us *α* + 2(*w* – 1) k-mers in total, with their relative orders corresponding to a permutation in *S*_*α*+2(*w* – 1)_. See Figure 3 for a visual example.

**Fig. 3:**
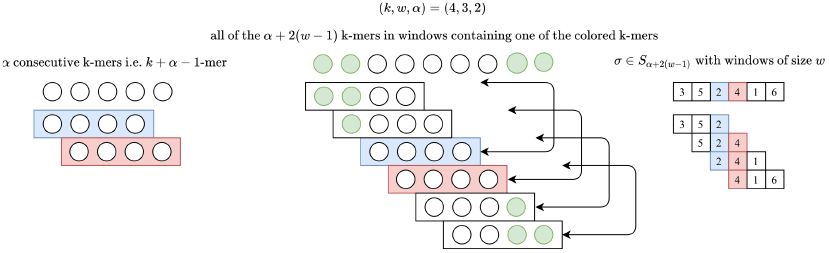
A graphical example of how *α* consecutive k-mers gives rise to a permutation in *S*_*α*+2(*w* – 1)_. The green bases correspond to extraneous bases, the colored k-mers correspond to the original *α* k-mers, and the colors/numbers in the permutation correspond to the k-mer and its relative order.

We proceed by counting permutations that correspond to some k-mer being chosen by a random minimizer method. We define a function *M*(*n, w, α, p*) which counts permutations in *S_n_* satisfying a general condition first.

#### Theorem 7

(Successful permutations for random minimizers). *Given parameters* (*n, w,α,p*) *with p* + *α* – 1 ≤ *n, let M*(*n, w,α,p*) *be the number of permutations in S_n_ such that for some window* [*σ*(*i*), …,*σ*(*i* + *w* – 1)], *the smallest element is one of σ*(*p*), *σ*(*p* + 1),…, *σ*(*p* + *α* – 1). *Then*

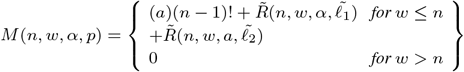

*where* 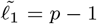, 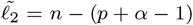 *and using* (*x*)_*n*_ *to mean the falling factorial*,

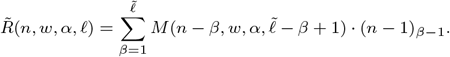

The specific choice of parameters (*n, w, α, p*) = (2(*w* – 1) + *α, w, α, w*) corresponds to the number of successful permutations for the minimizer given *α* consecutive k-mers since the *p* parameter describes the leftmost position of the first unmutated k-mer covering position *i*. Therefore, as before, Pr(*f, α*) = *M*(2(*w* – 1) + *α, w, α, w*)/(2(*w* – 1) + *α*)!. This theorem is proved Supplementary Section S3.

#### 4.2.1 Context dependency

The exact equality for Theorem 2 holds only when *f* is a 1-local method, which minimizers are not. Recall the exact expression using for calculating Cons(*f, θ, k*) in Equation 1. Consider the case

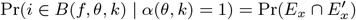

for *w* = 2 when *f* is a random minimizer. For *E_x_*, we must consider the 2(*w* – 1) + *α* = 3 relevant k-mers on *S*; call these k-mers *x*_1_, *x*_2_, *x*_3_, where *x*_2_ is the unmutated k-mer overlapping position *i*. We can see that Pr(*E_x_*) = 2/3 either from recognizing that this is the density 2/(*w* + 1) or from realizing that this is equivalent to permutations in S_3_ for which *σ*(2) = 1 or 2, where *σ*(2) is the relative ordering of *x*_2_.

However, the only k-mer on *S*′ which is guaranteed to be unmutated is *x*_2_, since *α* = 1. The surrounding k-mers 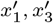 may not be equal to *x*_1_, *x*_3_. If they are not, then

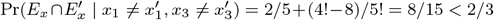

by considering a random ordering on 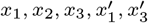 and seeing that if *x*_2_ is the smallest or second smallest, it is selected on both strings, but if it is the third smallest then there are 8 permutations for which *x*_2_ is not selected on one of the strings. Although 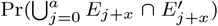 can be similarly phrased in terms of permutations, a trickier combinatorial problem arises. We take the route of analyzing the discrepancy in the upper bound of Theorem 2 empirically instead.

### 4.3 (*a, b, n*)-words method

We can also derive the probability vector for the previously (*a, b, n*)-words method which selects k-mers based on their prefix. We prove this result in Supplementary Section S4.

#### Theorem 8.

Pr(*f, α*) *under the* (*a, b, n*)-*words method is*

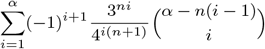

*where* 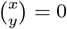 *if x* < 0.

## 5 Empirical results

We perform two sets of experiments. In Section 5.1 we compare Cons(*f, θ, k*) analytically and through simulations for a wider range of methods compared to previous studies (Frith *et al.*, 2020; Edgar, 2021) which focused only on simple minimizer methods and variations on their own methods. In Section 5.2 we modify the minimap2 (Li, 2018) software to use open syncmers and include experiments showing that alignment sensitivity is increased. While previous studies have focused on wholegenome alignment (Frith *et al.*, 2020; Edgar, 2021) as a use case for better conserved selection methods, we focus on long-read alignment as a possible use case.

### 5.1 Comparing Cons(*f, θ, k*) across different methods

In Section 4 we derived Pr(*f*) for four methods, and since three of those are 1-local methods we can calculate Cons(*f,θ,k*) in closed-form for these three methods using Theorem 2. In this section, we empirically calculate Cons(*f, θ, k*) for three more methods mentioned in Section 2.2: the random minimizer (for which we know Pr(*f*), giving an upper bound), the miniception, and a words based method with a highly optimized choice of W. Fixing some density *d,* we compare Cons(*f, θ, k*) to *UB*(*d*) · Pr(*α*(*θ, k*)), where *UB*(*d*) is defined in Section 3.1 as the upper bound on probability vectors. For the miniception method, it is not obvious how parameters affect *d* so we let *k*_0_ = *k* – *w* for *d* = 2/(*w* + 1) as suggested in Zheng *et al.* (2020a), and then modify *k*_0_ and *w* slightly until we get the density close to *d*. For the words method, we use two choices of *W* as *W*_4_ and *W*_8_, where the corresponding methods for *W*_4_ has density 1/4 and *W*_8_ has density 1/8. These sets are empirically found to perform well in Frith *et al.* (2020). We define these sets in Supplementary Section S6.

For methods with a closed-form for Cons(*f, θ, k*), we plot the exact value. For methods without a closed-form, we ran 100 simulations with |*S*| = 50, 000. We report mean and 95% confidence intervals (assuming normality) of Cons(*f, θ, k*) for these methods. Code can be found at https://github.com/bluenote-1577/local-kmer-selection-results.

We first fix *d* = 1/4, *k* = 17 and plot the fraction of upper bound achieved, 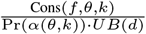, for all methods over a range of *θ*. We then fix *d* = 1/8, *k* = 25 and do a similar plot. For the (*a, b, n*)-words method, the closest choice of parameters leading to the most similar density is *n* = 2, 3 which gives density = 9/64 ~ 1/7.11 and 27/256 ~ 9.48; we plot both of these for reference. Both results are shown in Figure 4.

**Fig. 4:**
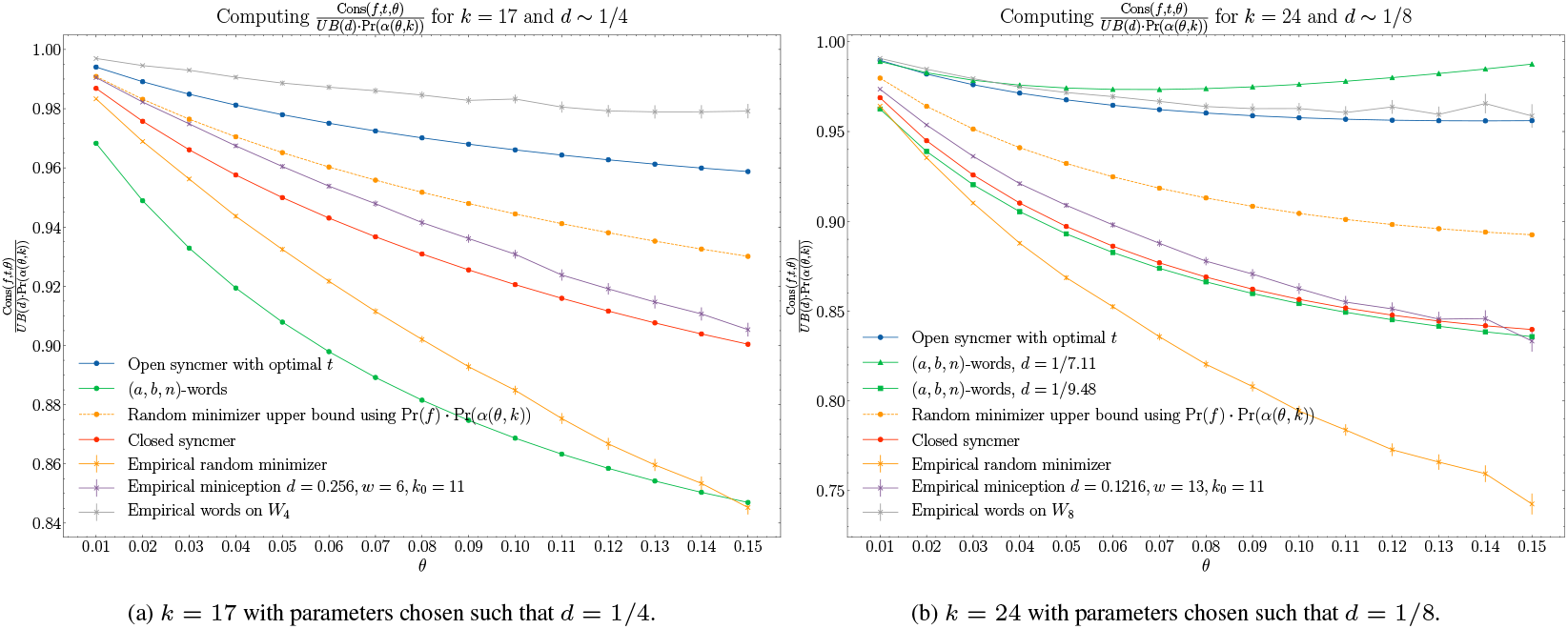
Fraction of upper bound achieved for conservation. 95% confidence intervals are built from 100 simulations for methods with empirically deduced values. Note that some methods have different densities due to parameter constraints; this is mentioned in the labels.

The results in Figure 4 show that the words method based on the set *W*_4_, *W*_8_ and the open syncmer methods perform well compared to the other methods. The large drop in conservation between the empirical random minimizer and the upper bound for the random minimizer indicates that context dependency plays a highly non-trivial role in conservation. Interestingly, the miniception algorithm performs reasonably well, even outperforming the 1-local closed-syncmer method despite suffering from the same context dependency issue. This is because of the design of the miniception which uses 1-local closed syncmers as “seeding” k-mers.

These results also tell us that the best methods already achieve ≥ 0.96 fraction of the possible upper bound *UB*(*d*) · Pr(*α*(*θ, k*)) for reasonable parameters and error rates, showing that there is not room for drastic improvement.

### 5.2 Using open syncmers in minimap2

Based on results from the previous section, we deduce that open syncmers and words based methods with specially chosen *W* perform the best with respect to conservation. We now ask how this result transfers to applications, particularly, in the setting of read mapping. To address this, we modified minimap2 (Li, 2018), a state-of-the-art read aligner, so that open syncmers are used instead of minimizers. We decided on using open syncmers because although its conservation is slightly less compared to words based methods, it allows the user to choose a range of densities without constructing a new words set *W* for each density.

Minimap2 aligns reads using a seed-chain-extend procedure. Roughly speaking, this works by firstly applying a k-mer selection method to all reads and reference genomes. Selected k-mers on the reads are then used as *seeds* to be matched onto the selected k-mers of the reference. Colinear sets of k-mer matches are collected into *chains,* and then dynamic programming based alignment is performed to fill gaps between chains. Our modification was to swap out the k-mer selection method, originally random minimizers, to an open syncmer method instead. For the rest of this section, use *k* = 15, *w* = 9 when using random minimizers and *k* = 15, *s* = 11 when using open syncmers so that the theoretical density of both methods are fixed at *d* =1/5. Our version of minimap2 can be found at https://github.com/bluenote-1577/os-minimap2.

We operate on three sets of real and simulated publicly available long-read datasets:

1. Real microbial PacBio Sequel long-reads from PacBio (2019), available at https://tinyurl.com/uhuwvxb8 and their corresponding assemblies.
2. Real human ONT nanopore reads from Miga *et al.* (2020) available at https://github.com/marbl/CHM13 (id: rel3; downsampled and only including reads > 1 kb in length) and the corresponding assembly CHM13.
3. Simulated PacBio long-reads on human reference GRCh38 using PBSIM(Ono *et al.*, 2013).

#### 5.2.1 Chaining score improvement

For each alignment of a read, minimap2 computes a *chaining score*, the exact details of which can be found in Section 2.1 in Li (2018). This chaining score measures the goodness of the best possible chain for that alignment. Roughly speaking, if k-mers in a chain overlap and do not have gaps, the chaining score is high. We took the long-read sequences and assembly for E. coli W (bc1087) in the PacBio (2019) dataset and compared the mappings for open syncmers versus minimizers. This is shown in Figure 5.

**Fig. 5:**
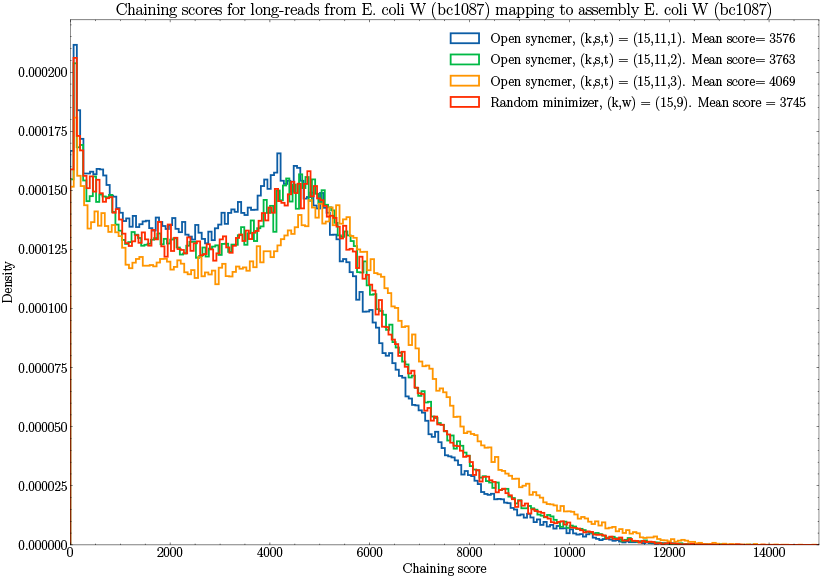
Histogram of chaining scores corresponding to alignments of reads for E. coli W (bc1087) in PacBio (2019) against its assembly. Minimap2 with open syncmers and minimizers were compared against each other with parameters chosen so that density is fixed at 1/5. Mean chaining scores are given in the legend. *t* =3 is the optimal value of t by Theorem 6.

For *t* = 3, the optimal parameter of *t*, open syncmers leads to better chaining scores. The theoretical value of Cons(*f, θ,k*) increases as *t* increases for open syncmer methods because the vectors Pr(*f*) are increasing component-wise as *t* increases; see Supplementary Figure 2. Therefore the increase in mean chaining score as t increases is suggestive that goodness of chaining is indeed correlated with conservation, even for PacBio reads which have higher rates of indels than substitutions (Dohm *et al.*, 2020). This suggests that the results from the conservation framework is still valid under realistic mutation models.

It is well known that empirical densities may deviate from theoretical densities for minimizers (Marçais *et al.*, 2017). To verify that the increase in chaining was not because of open syncmers having higher density than minimizers, we calculated the empirical density for minimizers/open syncmers on the reference genome in this experiment and found that it was 1/4.867 and 1 /5.059 respectively. This shows that even though the density for open syncmers was *lower* than for minimizers, the chaining score was higher.

#### 5.2.2 Alignment sensitivity and specificity analysis

We now examine how alignment sensitivity increases when using a more conserved k-mer selection method. We fix *t* = 3 with the same parameters for *k, w, s* as above. For the real datasets we analyze mapping quality and number of mapped reads. The difference between conservation of k-mer selection methods is more pronounced when the rate of mutation is higher, so we test how alignment changes as the reference genomes diverge from the reads. We test this effect by aligning the reads to varying reference genomes as well. The results are summarized in Table 3.

**Table 3.** We fix parameters (*k, s, t, w*) = (15, 11, 3, 9). Let *R*′ be the set of reads that were unmapped by minimap2 onto the reference for one or both of the methods. *OS* ⊂ *R*′ is the subset of reads that was successfully mapped using open syncmers, and *M* ⊂ *R*′ similarly for minimizers. 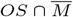 is the set of reads which are uniquely mapped by open syncmers, and 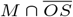 are reads uniquely mapped by minimizers. The average mapQ outputted by minimap2 within the set is presented as well. The datasets are described in Section 5.2.

| Long-read dataset | Reference | $\vert OS \cap \overline{M}\vert $ (mapQ) | $\vert M \cap \overline{OS}\vert $ (mapQ) | $\vert R'\vert $ | Total # reads |
| --- | --- | --- | --- | --- | --- |
| E. coli W (bc1087) | E. coli W (bc1087) | 312 (21.56) | 102 (11.57) | 2758 | 196901 |
| E. coli K12 (bc1106) | E. coli W (bc1087) | 548 (20.30) | 187 (11.33) | 6995 | 226906 |
| K. Pneumoniae (bc1074) | E. coli W (bc1087) | 11434 (19.80) | 3679 (12.10) | 119548 | 251838 |
| Downsampled human ONT (rel3) | Human - CHM13 | 370 (3.53) | 103 (2.52) | 13761 | 51210 |
| Downsampled human ONT (rel3) | Mouse - GRCm38 | 2467 (2.32) | 1005 (1.90) | 34463 | 51210 |

For the simulated reads, we used PBSIM Ono *et al.* (2013) to simulate PacBio CLR reads at 0.5 coverage across GRCh38 with mean length 15kb and 10% error rate. We report the mapping accuracy using paftools from Li (2018); a similar evaluation procedure is used in Jain *et al.* (2020). paftools calls an alignment correct if the overlap between an alignment and the true position of the simulated read is greater than a 0.1 fraction of all bases covering the true and mapped positions. We report accuracy and computational details in Table 4.

**Table 4.** Alignment of PBSIM simulated PacBio reads on human genome GRCh38 with 103630 reads, mean length 15kb, and 10% error rate. Parameters were chosen so that (*k, s, t, w*) = (15, 11, 3, 9). The number of unmapped reads was negligible. % of unique selected k-mers refers to k-mers selected by syncmers/minimizers on the reference genome that only appear once.

| Selection Method | % Incorrectly mapped reads | Indexing time (s) | Mapping time (s) | % of selected k-mers which are unique | Density |
| --- | --- | --- | --- | --- | --- |
| Open syncmer | 2.96% | 109 | 1150 | 27.9% | 0.1867 |
| Random minimizer | 2.89% | 95 | 979 | 37.0% | 0.1967 |

These results show that the number of reads that were uniquely mapped using open syncmers is consistently greater than with minimizers. In the case of human ONT reads mapping to the mouse genome, the relative increase in number of mapped reads is 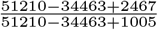 which is about an 8.2% increase. This effect increases as the reads and reference genomes become more divergent and mapping becomes more challenging. There is a slightly worse accuracy for the open syncmer method, which is to be expected. An important observation is that the open syncmers tend to select non-unique k-mers more often, which is not ideal. It would be interesting to analyze how different k-mer selection methods interact with the repetitive nature of genomes; for example, we used a random ordering for the syncmer construction, but perhaps a better ordering would select more unique k-mers.

## 6 Conclusion

In summary, we first described a new mathematical framework for understanding k-mer selection methods in the context of conservation. Our framework allowed us to prove several results pertaining to upper bounds, optimal parameter choices, and closed-form expressions for conservation. We then investigated conservation empirically and then found that augmenting minimap2 with a better conserved method increases alignment sensitivity.

There are some interesting theoretical and practical problems still to be addressed. A notable result of ours is that the best methods can already achieve > 0.96 of the upper bound for conservation for certain parameters, implying that major improvements using our framework are not possible. However, real genomes do not consist of i.i.d uniform letters. There are a mix of high-complexity and low-complexity regions in genomes, so techniques for better distributed selection k-mers can be investigated (Jain *et al.*, 2020) for example. Sequence-specific k-mer selection methods, where the selection method is specifically tuned for a certain string is another area of practical importance (Zheng *et al.*, 2021). Theoretical problems we think would be interesting include understanding how tight the bound given in Section 3.1 is when parameters are not in the asymptotic regime and deeper analysis on the context dependency problem for random minimizers.

An orthogonally related recent idea are strobemers (Sahlin, 2021), which have been proposed as a k-mer alternative for sequence mapping. It has been shown that strobemers allow for much higher conservation (called match-coverage in Sahlin (2021)) than k-mers. Understanding selection techniques for k-mer replacements is an area of unexplored research where we believe that some of our techniques may be useful.

