## Supplementary Materials for "Theory of local k-mer selection with applications to long-read alignment"

### Contents

**S1** $\Pr(\alpha(\theta, k))$ **formula and figure** **1**

**S2 Open syncmer proofs** **2**

**S3 Proof of random minimizer probability vector** **4**

**S4 Proof of** $(a, b, m)$**-words method probability vector** **5**

**S5 Comparing** $\Pr(f)$ **7**

**S6 Defining** $W_4, W_8$ **7**

### S1 $\Pr(\alpha(\theta, k))$ formula and figure

**Theorem S1.1.** *For $2k-1$ i.i.d Bernoulli trials with success probabillity $1-\theta$ and $0 \leq \beta \leq k-1$,*

$$\Pr(\alpha(\theta, k) = \beta + 1) = \Pr(\textit{Longest run of successes is } k + \beta)$$
$$= \sum_{b=0}^{k-\beta-2} T_k(\beta, b)(1-\theta)^{k+\beta+b} \cdot \theta^{k-\beta-b-1}$$

*where*

$$T_k(\beta, b) = 2\binom{k-2-\beta}{b} + (k-\beta-2)\binom{k-3-\beta}{b}$$

*and binomial coefficients with negative parameters are 0. For $\beta = k-1$, the probability of $2k-1$ successes is just $(1-\theta)^{2k-1}$.*

*Proof.* Suppose $\beta < k-1$. If the maximum successful run is of length $k+\beta$ in $2k-1$ trials, this must be the *only run* of $k+\beta$ successes in a row. Label the start and end of this sequence by positions $i, j \in \{1, ..., 2k-1\}$, where $j = i+k+\beta-1$. The possible positions of $i$ are $i \in \{1, ...., k-\beta\}$. We calculate $\sum_{k=1}^{k-\beta} \Pr(k+\beta \text{ sucesses in a row}, i = k)$.

Case 1: if $i = 1$ or $i = k-\beta$, then trial $i+1$ or $i-1$ has to be a failure respectively, otherwise the run is longer than $k+\beta$. There are $2k-1-(k+\beta+1) = k-\beta-2$ remaining trials which can be either successes for failures. Letting $b$ be the number of successes in the rest of the trials and conditioning on $b$, we get the probability of $i = 1$ or $i = k-1$ as

$$2\sum_{b=0}^{k-\beta-2}\binom{k-2-\beta}{b}(1-\theta)^{k+\beta+b}\theta^{k-\beta-b-1}.$$

Case 2: if $i \neq 1$ and $i \neq k-\beta$, then both of the trials $i-1$ and $j+1$ have to be failures. This leaves us with $k-\beta-3$ remaining trials. Conditioning on $b$ again, we get the probability of $i = 2, ..., k-2$ as

$$(k-\beta-2)\sum_{b=0}^{k-\beta-3}\binom{k-\beta-3}{b}(1-\theta)^{k+\beta+b}\theta^{k-\beta-b-1}.$$

Summing the probabilities together yields the result when $\beta < k-1$. If $\beta = k-1$ then clearly the probability is just $(1-\theta)^{2k-1}$. □

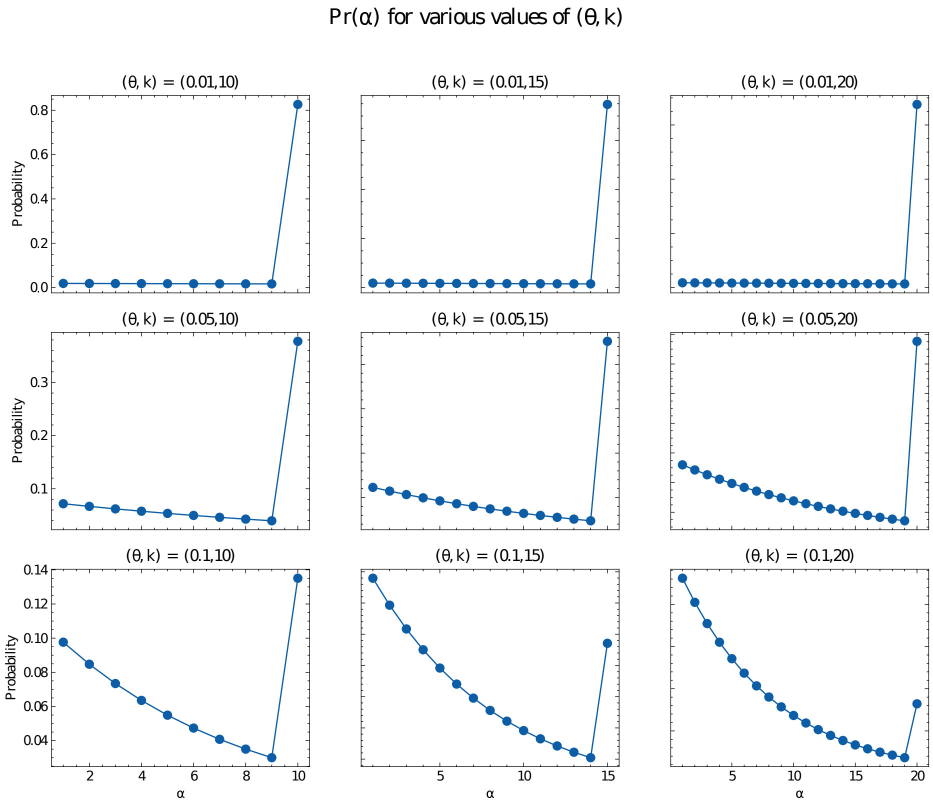


Figure 1: $\Pr(\alpha(\theta,k)) = [\Pr(\alpha(\theta,k)=1),...,\Pr(\alpha(\theta,k)=k)]$ for various values of $\theta$ and $k$.

### S2 Open syncmer proofs

**Theorem S2.1** (Successful permutations for open syncmers)**.** *Using parameters $k, s, t$ as defined in the definition of open syncmers let $\tau = t-1$ and $OS(\alpha,k,s,t)$ be the number of permutations in $S_{k-s+\alpha}$ such that for some window $[\sigma(i),...,\sigma(i+k-s)]$ the smallest element is $\sigma(i+\tau)$. Define $\ell_1 = \tau, \ell_2 = k-s-\tau$. Then*

$$OS(\alpha,k,s,t) = \alpha(k-s+\alpha-1)! + R(\alpha,k,s,t,\ell_1) + R(\alpha,k,s,t,\ell_2).$$

*We define $R(\alpha,k,s,t,\ell)$ as*

$$R(\alpha,k,s,t,\ell) = \sum_{\beta=1}^{\ell} (k-s+\alpha-1)_{\beta-1} OS(\alpha-\beta,k,s,t)$$

*where the subscript indicates falling factorial, and $OS(\alpha-\beta,k,s,t)=0$ if $\beta \geq \alpha$.*

This is proved in the Appendix.

*Proof.* We condition on the position of the smallest element, i.e. the index $\beta$ for which $\sigma(\beta)=1$. Let the set $A_\tau = \{\tau+1, \tau+2, ..., \tau+\alpha\}$

**(Case 1 - if $\beta \in A_\tau$).** In this case, the window

$$[\sigma(\beta-\tau),...,\sigma(\beta-\tau+(k-s))]$$

is valid and has the desired property that $\sigma((\beta + -\tau) + (\tau)) = \sigma(\beta)$ is the smallest integer in the window, so these permutations all satisfy condition 2 above. There are $\alpha(k - s + \alpha - 1)!$ such permutations.

**(Case 2 - if** $\beta < \tau + 1$**).** In this case, $\beta$ is left of position $t$. Notice that for all windows containing position $\beta$ will never be successful since the first window contains $\beta$ at position $< \tau + 1$, and the relative position of $\beta$ in subsequent windows will be $< \tau + 1$ as well.

The remaining windows which may still satisfy condition 2 lie are sub-windows of $[\sigma(\beta+1)...\sigma(k-s+\alpha)]$, which may be considered a permutation in $S_{k-s+\alpha-\beta}$ after relabelling elements to be in $\{1, ..., k - s + \alpha - \beta\}$ to preserve the relative order.

This new permutation has to satisfy condition 2, and the number of such permutations is exactly $OS(\alpha - \beta, k, s, t)$. We have to multiply by an additional $(k - s + \alpha - 1)_{\beta-1}$ to count the possible values for the $\beta - 1$ entries to the left of $\beta$, each of which give the same permutation in $S_{w+a-b}$ after relabelling. Summing over $b = 1, ..., \tau = \ell_1$ gives the $R(\alpha, k, s, t, \ell_1)$ term.

**(Case 3 - if** $\beta > \tau + \alpha$**).** This case is identical to case 2 and the same argument works after flipping directions. This works by summing over the $\ell_2 = k - s - \tau$ possible positions $\beta \in \{k - s + \alpha, k - s + \alpha - 1, ..., \tau + 1 + \alpha\}$ and using a the same relabelling after cutting off a portion of the permutation. The number of permutations for $\beta = k - s + \alpha - i$ is the same as for $\beta = i$ by symmetry. Using this correspondence gives the $R(\alpha, k, s, t, \ell_2)$ term and completes the proof.

□

We now prove the following theorem.

**Theorem S2.2.** *Let* $\hat{t} = \lceil \frac{k-s+1}{2} \rceil$*. Then* $OS(\alpha, k, s, \hat{t}) \geq OS(\alpha, k, s, t)$ *for any valid choice of* $t$*.*

**Lemma S2.3.** *Fix* $k, s, t, \alpha$ *and define* $(k - s + \alpha - \beta - 1)_{\beta-1} OS(\alpha - \beta, k, s, t) = \overline{OS}(\alpha, \beta, t)$*. If* $\gamma \geq \beta$*, for any* $t$*, we have*

$$\overline{OS}(\alpha, \beta, t) \geq \overline{OS}(\alpha, \gamma, t).$$

*Proof of Lemma.* We show $\overline{OS}(\alpha, \beta - 1, t) \geq \overline{OS}(\alpha, \beta, t)$ for any $\beta$, which implies the result. This is equivalent to showing that

$$OS(\alpha - \beta + 1, k, s, t) \geq \frac{(k - s + \alpha - 1)_{\beta-1}}{(k - s + \alpha - 1)_{\beta-2}} OS(\alpha - \beta, k, s, t)$$

$$= (k - s + \alpha - \beta + 1) OS(\alpha - \beta, k, s, t).$$

Notice that

$$OS(\alpha - \beta + 1, k, s, t)/(k - s + \alpha - \beta + 1)! = \Pr(f, \alpha - \beta + 1)$$

and

$$OS(\alpha - \beta, k, s, t)/(k - s + \alpha - \beta)! = \Pr(f, \alpha - \beta)$$

when $f$ is an open syncmer method with fixed parameters $k, s, t$ from our correspondence between random permutations and the event a k-mer is selected by $f$. By definition, $\Pr(f, \alpha - \beta + 1) \geq \Pr(f, \alpha - \beta)$. Technically, the correspondence is only true up to a small error due to the chance of repeated k-mers appearing in a window, but one can make $OS(x, k, s, t)$ arbitrarily close to $\Pr(f, x)$ by letting the alphabet be very large, making repeats unlikely (see the Section 2.3.1 in [1]). Then

$$OS(\alpha - \beta + 1, k, s, t) \geq (k - s + \alpha - \beta + 1) OS(\alpha - \beta, k, s, t)$$

follows from $\Pr(f, \alpha - \beta + 1) \geq \Pr(f, \alpha - \beta)$, and we're done.

□

*Proof of Theorem S2.2.* We use the similar notation as Lemma S2.3 for $\overline{OS}$.

Observe that

$$OS(\alpha, k, s, t) = OS(\alpha, k, s, k - s + 2 - t)$$

since this just swaps the $\ell_1, \ell_2$ in the definition. Since $k - s + 2 - \hat{t} = \hat{t}$ or $\hat{t} + 1$ depending on if $k - s + 1$ is odd or even, we only need to prove that this inequality holds for $t < \hat{t}$. We will assume $k - s + 1$ is odd for exposition since the indices are easier to handle, but the result holds either way after a slight modification.

We proceed by induction on $\alpha$ for $OS(\alpha, k, s, t)$. For the base case $\alpha = 1$, notice that clearly $OS(1, k, s, t) = OS(1, k, s, \hat{t}) = (k - s)!$. Let $\hat{\tau} = \hat{t} - 1$ and $\tau = t - 1$. Therefore we want the following term to be positive:

$$\begin{aligned} OS(\alpha, k, s, \hat{t}) - OS(\alpha, k, s, t) = & \sum_{\beta=1}^{\tau} [\overline{OS}(\alpha, \beta, \hat{t}) - \overline{OS}(\alpha, \beta, t)] \\ & + \sum_{\beta=1}^{k-s-\hat{\tau}} [\overline{OS}(\alpha, \beta, \hat{t}) - \overline{OS}(\alpha, \beta, t)] \\ & + \sum_{\beta=\tau+1}^{\hat{\tau}} \overline{OS}(\alpha, \beta, \hat{t}) - \sum_{\beta=k-s-\hat{\tau}+1}^{k-s-\tau} \overline{OS}(\alpha, \beta, t) \end{aligned} \tag{1}$$

By the induction assumption the first two sums are $\geq 0$ since $OS(\alpha - \beta, k, s, \hat{t}) \geq OS(\alpha - \beta, k, s, t)$ for $\beta \geq 1$. For the last term, $k - s + 1$ odd gives us $k - s - \hat{\tau} = \hat{\tau}$. We can rewrite the last line as

$$\sum_{j=1}^{\hat{\tau}-\tau} \overline{OS}(\alpha, (\hat{\tau} - j + 1), \hat{t}) - \overline{OS}(\alpha, (\hat{\tau} + j), t).$$

By the induction assumption, $\overline{OS}(\alpha, (\hat{\tau} - j + 1), \hat{t}) \geq \overline{OS}(\alpha, (\hat{\tau} - j + 1), t)$ and using Lemma S2.3 finishes the proof because $\hat{\tau} - j + 1 \leq \hat{\tau} + j$ for all $j \geq 1$.

□

We show in Figure 2 what $\Pr(f)$ looks like for syncmer methods over a range of $t$.

### S3 Proof of random minimizer probability vector

**Theorem S3.1** (Successful permutations for random minimizers)**.** *Given parameters* $(n, w, \alpha, p)$ *with* $p + \alpha - 1 \leq n$*, let* $M(n, w, \alpha, p)$ *be the number of permutations in* $S_n$ *such that for some window* $[\sigma(i), ..., \sigma(i + w - 1)]$*, the smallest element is one of* $\sigma(p), \sigma(p + 1), ..., \sigma(p + \alpha - 1)$*. Then*

$$M(n, w, \alpha, p) = \left\{ \begin{array}{ll} (a)(n-1)! + \tilde{R}(n, w, \alpha, \tilde{\ell_1}) + \tilde{R}(n, w, a, \tilde{\ell_2}) & \textit{for } w \leq n \\ 0 & \textit{for } w > n \end{array} \right\}$$

*where* $\tilde{\ell_1} = p - 1$*,* $\tilde{\ell_2} = n - (p + \alpha - 1)$ *and using* $(x)_n$ *to mean the falling factorial,*

$$\tilde{R}(n, w, \alpha, \ell) = \sum_{\beta=1}^{\tilde{\ell}} M(n - \beta, w, \alpha, \tilde{\ell} - \beta + 1) \cdot (n-1)_{\beta-1}.$$

*Proof.* As in the proof of Theorem S2.1, we condition on the position of the smallest element, i.e. the index $\beta$ for which $\sigma(\beta) = 1$. Let the set $A_p = \{p, p + 1, ..., p + \alpha - 1\}$.

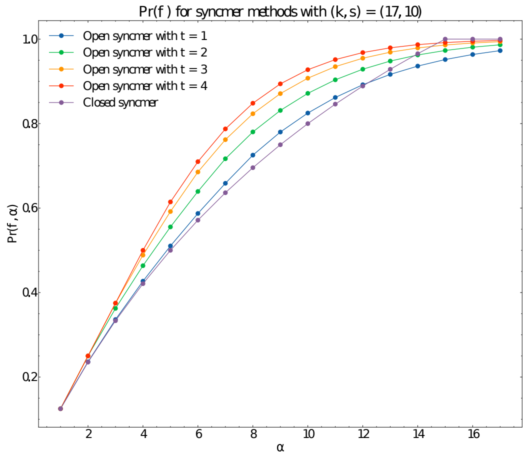


Figure 2: The probability vector for open syncmers with varying $t$ parameters $(k, s) = (17, 10)$. The $s$ parameter for the closed syncmer was chosen so that the densities are equal. We only evaluate for $t \leq 4$ because $t \mapsto k - s + 2 - t$ gives the same probabilities for open syncmers by Theorem S2.1.

**(Case 1 - if $\beta \in A_p$).** This permutation clearly is successful. There are $\alpha(n-1)!$ such permutations.

**(Case 2 - if $\beta < p$).** In this case, $\beta$ is left of position $p$. All windows containing position $\beta$ will never be successful since $\beta \notin A_p$ and $\sigma(\beta)$ is the smallest element in the window. The only possible successful windows are the sub-windows of $[\sigma(\beta+1)...\sigma(n)]$. We can relabel the positions after shifting by $\beta$ and consider this as a new permutation on $1, ..., n-\beta$ after relabelling $\sigma(i)$ while preserving relative order. This sub-problem is exactly counted by $(n-1)_{\beta-1}M(n-\beta, w, \alpha, p-\beta)$ after multiplying by the $(n-1)_{\beta-1}$ possible values for $\sigma(i)$, $i < \beta$. Notice that if $n - \beta < w$, then there are no windows that satisfy our requirement, so $M(n-\beta, w, a, p-\beta) = 0$. Summing over $\beta < p$ gives

$$\sum_{\beta=1}^{p-1} (n-1)_{\beta-1} M(n-\beta, w, \alpha, p-\beta).$$

**(Case 3 - if $\beta > p + \alpha - 1$).** The exactly same argument follows as in case 2. We see that successful windows must be sub-windows of $[\sigma(1), ..., \sigma(\beta-1)]$, so this is almost counted by $(n-1)_{n-\beta}M(\beta-1, w, \alpha, p)$ over all $\beta > p + \alpha - 1$. We can shift indices to get

$$\sum_{\beta=p+\alpha}^{n} (n-1)_{n-\beta} M(\beta-1, w, \alpha, p) = \sum_{\beta=1}^{n-(p+\alpha-1)} M(n-\beta, w, \alpha, p)(n-1)_{\beta-1}.$$

To rewrite the equation to be in a similar form to case 2, one can see that $M(n-\beta, w, \alpha, p) = M(n-\beta, w, \alpha, (n-\beta) - (p+\alpha-1) + 1)$ which corresponds to "flipping' the permutation on $S_{n-\beta}$ so that position $i \mapsto n - \beta - i + 1$. This completes the proof. □

### S4 Proof of $(a, b, m)$-words method probability vector

**Theorem S4.1.** $\Pr(f, \alpha - 1)$ *under the* $(a, b, n)$*-words method is*

$$\sum_{i=1}^{\alpha}(-1)^{i+1}\frac{3^{ni}}{4^{i(n+1)}}\binom{\alpha-n(i-1)}{i}$$

*where* $\binom{x}{y}=0$ *if* $x<0$.

We first prove an intermediate combinatorial lemma.

**Lemma S4.2.** *Given a set of of* $\alpha$ *elements labelled* $\{1,...,\alpha\}$*, the number of ways* $c(n+1,i)$ *to choose* $i$ *elements* $x_1,...,x_i$ *where we order* $x_j<x_{j+1}$ *for* $j=1,...,i-1$ *and* $|x_j-x_{j+1}|\geq(n+1)$ *for all* $j$ *is*

$$\binom{\alpha-n(i-1)}{i}.$$

*Proof.* Let $y_0=x_1-1$, $y_1=x_2-x_1-1,...,y_i=\alpha-x_i$. The $y_i$s represent the gaps between $x_i$s and also the endpoints. A valid choice of $x_i$s corresponds exactly to a choice of $y_i$s such that each $y_i\geq n$ for $i=1,...,i-1$ and $y_0,y_i\geq 0$. Furthermore,

$$\sum_{j=0}^{i}y_i=\sum_{j=1}^{i-1}(x_{i+1}-x_i-1)+\alpha-x_i+x_1-1=\alpha-i$$

We can take $z_j=y_j-n$ for $i=1,...i-1$ and $z_j=y_j$ otherwise to get the equivalent problem of finding $z_j$ all $\geq 0$ such that

$$\sum_{j=0}^{i}z_j=\alpha-i-(i-1)n.$$

This problem is equivalent to putting $\alpha-i-(i-1)n$ indistinct balls into $i+1$ distinct jars represented by the variables $z_j$. The solution is

$$\binom{[i+1]+[\alpha-i-(i-1)n]-1}{\alpha-i-(i-1)n}=\binom{\alpha-n(i-1)}{i}$$

as desired.

□

*Proof of Theorem S4.1.* The probability that at least one of the k-mers is selected is

$$\Pr(\bigcup_{i=1}^{\alpha}E_i)$$

where $E_i$ is the event that the $i-$th k-mer is selected. By inclusion-exclusion, we get

$$\Pr(\bigcup_{i=1}^{\alpha}E_i)=\sum_{I\subset\{1,...,\alpha\}}(-1)^{|I|+1}\Pr(E_I)=\sum_{i=1}^{\alpha}(-1)^{i+1}\sum_{I\subset\{1,...,\alpha\},|I|=i}\Pr(E_I)$$

where $E_I=\bigcup_{i\in I}E_i$. Now note that the probability that $E_\alpha\cap E_\beta$ for $|\alpha-\beta|<n$ occurs is 0; k-mers with prefix *abbb...* may not be within distance $n+1$ from each other. If the $i$ k-mers are all distance $\geq n+1$ apart, then the probability of that event occurring is just $(\frac{3^n}{4\cdot 4^n})^i$ because this is just the sequence *abbb...* appearing $i$ times in a string of i.i.d random letters. Therefore, denoting $c(i,n+1)$ to be the number of ways to select $i$ elements from $\{1,...,\alpha\}$ such that each element is at least pairwise distance $n+1$ apart, we get

$$\sum_{i=1}^{\alpha}(-1)^{i+1}\sum_{I\subset\{1,...,\alpha\},|I|=i}\Pr(E_I)=\sum_{i=1}^{\alpha}(-1)^{i+1}c(n+1,i)\frac{3^{ni}}{(4^{n+1})^i}.$$

Plugging in the above lemma finishes the proof.

□

### S5 Comparing $\Pr(f)$

In Figure 3, we plot all $\Pr(f)$ and $UB(d)$ where all methods have density $d = 1/7$ except for the words method, which has density $9/64 \sim 1/7.11$. This is due to the limited range of parameters choices for the methods.

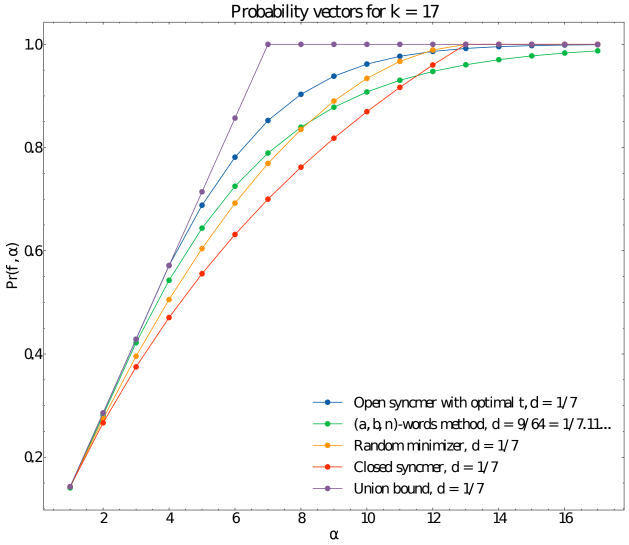


Figure 3: Comparison of $\Pr(f)$ for all methods with exact distributions derived. Note that the density for the words method is slightly smaller.

### S6 Defining $W_4, W_8$

We take the words set $W_4$ as

$$\begin{aligned} W_4 = \{&rrrrry, rryrry, rryryy, ryrrrr, \\ &ryrrry, ryryry, ryyrrr, ryyrry, \\ &ryyryr, ryyryy, ryyyry, ryyyyy, \\ &ryrrry, ryryry, ryyrrr, ryyrry\}. \end{aligned} \tag{2}$$

Here $r = \{A, G\}$ and $y = \{C, T\}$ and we mean $rryrry$ to be all 6-mers that satisfy this condition. This set leads to a $d = 1/4$ method, and was found by an optimization algorithm [2].

We take the words set $W_8$ as

$$\begin{aligned} W_8 = \{&rrrrrrry, rryrrryy, ryrrrryr, ryrrrryy, \\ &ryrrryry, yrrrrrry, yrrrrryr, yrrrrryy, \\ &yrrryrry, yrryrryr, yrryrryy, yryrrryy, \\ &yryrryry, yryrryyr, yryrryyy, yryryryy, \\ &yyrrrryr, yyrrrryy, yyrrryry, yyrrryyr, \\ &yyrrryyy, yyrryryr, yyrryryy, yyrryyry, \\ &yyrryyyr, yyrryyyy, yyryryyr, yyryryyy, \\ &yyryyryy, yyyryyyr, yyyryyyy, yyyyyyyr\} \end{aligned} \tag{3}$$

Here $r = \{A, G\}$ and $y = \{C, T\}$ and we mean *rryrry* to be all 6-mers that satisfy this condition. This set leads to a $d = 1/8$ method, and was found by an optimization algorithm [2].
